# Explication of CB_1_ receptor contributions to the hypothermic effects of Δ9-tetrahydrocannabinol (THC) when delivered by vapor inhalation or parenteral injection in rats

**DOI:** 10.1101/2020.05.22.111468

**Authors:** Jacques D. Nguyen, K. M. Creehan, Yanabel Grant, Sophia A. Vandewater, Tony M. Kerr, Michael A. Taffe

## Abstract

The use of Δ^9^-tetrahydrocannabinol (THC) by inhalation using e-cigarette technology grows increasingly popular for medical and recreational purposes. This has led to development of e-cigarette based techniques to study the delivery of THC by inhalation in laboratory rodents. Inhaled THC reliably produces hypothermic and antinociceptive effects in rats, similar to effects of parenteral injection of THC. This study was conducted to determine the extent to which the hypothermic response depends on interactions with the CB_1_ receptor, using pharmacological antagonist (SR141716, AM-251) approaches. Groups of rats were implanted with radiotelemetry devices capable of reporting activity and body temperature, which were assessed after THC inhalation or injection. SR141716 (4 mg/kg, i.p.) blocked or attenuated antinociceptive effects of acute THC inhalation in male and female rats. SR141716 was unable to block the initial hypothermia caused by THC inhalation, but temperature was restored to normal more quickly. Alterations in antagonist pre-treatment time, dose and the use of a rat strain with less sensitivity to THC-induced hypothermia did not change this pattern. Pre-treatment with SR141716 (4 mg/kg, i.p.) blocked hypothermia induced by i.v. THC and reversed hypothermia when administered 45 or 90 minutes after THC (i.p.). SR141716 and AM-251 (4 mg/kg, i.p.) sped recovery from, but did not block, hypothermia caused by vapor THC in female rats made tolerant by prior repeated THC vapor inhalation. The CB_2_ antagonist AM-630, had no effect. These results suggest that hypothermia consequent to THC inhalation is induced by other mechanisms in addition to CB_1_ receptor activation.

## 1. Introduction

A substantial and sustained decrease in body temperature, and a reduction in sensitivity to a noxious stimulus (antinociception), are two major indicators of cannabinoid activity in laboratory rodents (Abel, 1973; Borgen et al., 1973; Lichtman and Martin, 1990; Taylor and Fennessy, 1977; Vidal et al., 1984; Wiley et al., 2014). The exogenous cannabinoid compound Δ^9^-tetrahydrocannabinol (THC) also decreases the body temperature of monkeys (McMahon et al., 2005; Taffe, 2012), dogs (Hardman et al., 1971), cats (Schmeling and Hosko, 1980) and other species, as summarized (Clark and Clark, 1981). Curiously, data on temperature effects in humans are scarce but at least one case report indicates profound hypothermia in a child after cannabis resin ingestion (Bro et al., 1975) and another study indicates a small, but consistent, decrease in temperature after THC was ingested (Waskow et al., 1970). Some of the lack of clear evidence for thermoregulatory effects of THC in humans could be due to tolerance and potentially a limited range of dosing that has been employed. Human laboratory studies tend to be conducted in experienced cannabis users who may have become tolerant to any thermoregulatory effects, as is seen in preclinical models (Ginsburg et al., 2014; Nguyen et al., 2020).

Effects of THC on temperature appear to mostly be mediated via the CB_1_ receptor since the CB_1_ antagonist/inverse agonists SR141716 (Rimonabant) and AM-251 have been shown to block effects of cannabinoid agonists such as Δ^9^-tetrahydrocannabinol (THC) when delivered by parenteral injection (Grim et al., 2017; Taffe et al., 2015) and deletion of the CB_1_ receptor in mice prevents hypothermic responses to THC (Wang et al., 2019; Zimmer et al., 1999). Prior studies from this laboratory found that SR141716, administered 15 minutes prior to intraperitoneal injection of THC blocked the body temperature decreases, and tail-flick latency increases (Nguyen et al., 2016b; Taffe et al., 2015; Taffe et al., 2020), which is consistent with prior reports that SR141716 pretreatment attenuated a prolactin response to THC (Fernandez-Ruiz et al., 1997) and the hypothermia caused by the CB_1_ full agonist WIN 55,212-2 in mice (Son et al., 2010). In addition, SR141716 and another CB1 antagonist AM-251 prevented hypothermic and antinociceptive effects of THC, i.p., but failed to reverse hypolocomotor effects, in male C57BL/6J mice (McMahon and Koek, 2007). SR141716 only partially reversed rectal temperature hypothermia observed 2 h after injection in chair restrained rhesus monkeys (McMahon et al., 2005), but fully prevented the hypothermia in unrestrained monkeys for the first 3 hours after intramuscular injection in another study (Taffe, 2012); this difference may be due to the relative duration of action of SR141716 versus THC since the former paper did not report temperature prior to 2 h after injection. Additionally, SR141716 failed to reverse the effects of THC on operant responding in monkeys (McMahon et al., 2005).

The majority of human THC consumption occurs via the inhalation route from the smoking of combusted plant material or, more recently, inhalation of THC containing vapor created without combustion (Hazekamp et al., 2006; Morean et al., 2015; Morean et al., 2017; Perrine et al., 2019; Zuurman et al., 2008). Despite this, pre-clinical animal models of non-combusted cannabinoid inhalation have been relatively rare (Lichtman et al., 2000; Lichtman et al., 2001; Wilson et al., 2002) until recently. For example, it has been shown that intrapulmonary delivery of Δ^9^-tetrahydrocannabinol (THC), using a system based on commercial e-cigarette devices, results in a robust and dose-dependent hypothermia in adult (Javadi-Paydar et al., 2018; Nguyen et al., 2016b) and adolescent (Nguyen et al., 2020) rats of each sex, and a similar system results in hypothermia in mice following inhalation exposure to synthetic cannabinoid agonists (Lefever et al., 2017). This approach has also shown evidence for reward-seeking behavior in rats (Freels et al., 2020).

Antagonist effects of SR141716 have also been demonstrated following inhalation of THC (or cannabis smoke) but efficacy may be more variable. SR141716 blocked hypolocomotive effects of marijuana smoke inhalation in rats (Bruijnzeel et al., 2016) and fully blocked antinociception caused by aerosolized THC, but only partially blocked the antinociceptive effects of cannabis smoke inhalation, in male ICR mice (Lichtman et al., 2000; Lichtman et al., 2001). Catalepsy caused by marijuana smoke inhalation was not blocked by SR141716 in the latter study, but hypothermia was not produced by smoke inhalation. Interestingly, SR141716 fully reversed the antinociceptive, hypothermic and cataleptic effects of THC when administered *intravenously* in the same behavioral model (Lichtman et al., 2001). SR141716 blocked thermoregulatory, anti-nociceptive and locomotor effects of inhaled THC in Swiss Webster mice (Marshell et al., 2014) and our original paper showed that SR141716 blocked the antinociceptive effects of inhaled THC (Nguyen et al., 2016b), however, no evidence regarding the effect of SR141716 on THC inhalation-induced hypothermia was included in the latter study.

As was briefly reviewed above, SR141716 does not always fully block effects of THC and this may depend on outcome measure, species, THC dose or other as yet unexplicated factors such as the route of administration. This study was therefore designed to determine if the body temperature response to, and antinociceptive effects of, the inhalation of THC via e-cigarette type technology are attenuated by pre-treatment with CB_1_ antagonist/inverse agonists. As humans increasingly ingest THC via e-cigarette technology it has become increasingly important to develop translational pre-clinical models that are capable of evaluating the consequences of this route of administration. One important part of this development is determining similarities and differences with prior pre-clinical work with other routes of exposure.

The present study set out to determine if SR141716 is capable of blocking the hypothermia induced by the inhalation of THC vapor produced by e-cigarette devices. Our very first observation, conducted in the group of animals used for our initial report (Nguyen et al., 2016b), but not reported therein, found that SR141716 did not attenuate the initial drop in body temperature observed after THC inhalation. Body temperature did, however, appear to return to the normal range more quickly. More recently we have reported that SR141716 fails to block the hypothermia induced by the inhalation of THC in both Wistar and Sprague-Dawley male rats, despite attenuating the anti-nociceptive effects of inhaled THC, and preventing hypothermia caused by i.p. injection of THC, in each strain (Taffe et al., 2020). This study further explored these observations first by administering the SR141716 antagonist 90 minutes prior to the start of inhalation to match the apparent timing of the restoration of body temperature after the initial reduction. We next administered SR141716 after the cessation of inhalation (when body temperature was at ∼nadir) to determine if this would speed the restoration of normative temperature immediately, or after ∼90 minutes, thereby uncoupling the SR141716 and THC exposure latencies. Positive control experiments were next conducted to administer SR141716 45 minutes and 90 minutes after i.p. injection to further determine if SR141716 would exhibit efficacy in reversing a hypothermia during the initial, or well-engaged, stages of the response to i.p. injection of THC. We next administered a higher (10 mg/kg) dose of SR141716 in a group of Wistar rats. We’ve shown that Wistar are less sensitive to the temperature disrupting effects of THC compared with Sprague-Dawley rats (Taffe et al., 2020), thus the higher antagonist dose might be expected to confer even greater efficacy in a less-responsive rat strain. Finally we evaluated the effects of SR141716, another CB1 antagonist (AM-251) and the CB2 antagonist AM-630 in a group of female rats made tolerant to THC hypothermia via a twice-daily repeated THC regimen. It has been well established that repeated exposure to THC via parenteral injection produces tolerance to the acute effects in laboratory animals such as mice (Anderson et al., 1975; Fan et al., 1994), rats (Jarbe, 1978; Taylor and Fennessy, 1978; Wiley and Burston, 2014) or monkeys (Ginsburg et al., 2014; Smith et al., 1983; Winsauer et al., 2011). This tolerance is associated with a reduction in the efficacy of CB_1_ mechanisms thus, again, the hypothesis was that perhaps a given antagonist dose would exhibit greater efficacy. Finally, we evaluated plasma THC concentrations after vapor inhalation and i.p. injection of THC in male and female groups of Wistar and Sprague-Dawley rats to show that differences in SR141716 efficacy across routes of administration are not due to a substantially different systemic THC dose.

## 2. Methods

### 2.1. Subjects

Male (N=27) and female (N=29) Wistar (Charles River, New York) and male (N=22) and female (N=6) Sprague-Dawley (Harlan, Livermore, CA) rats were housed in humidity- and temperature-controlled (23±2 °C) vivaria on 12:12 hour light:dark cycles. Rats had *ad libitum* access to food and water in their home cages and all experiments were performed in the rats’ scotophase. Rats entered the laboratory at 10-11 weeks of age. All procedures were conducted under protocols approved by the Institutional Care and Use Committee of The Scripps Research Institute.

### 2.2. Radiotelemetry

Groups of female Wistar (N=16), male Sprague-Dawley (N=16) and male Wistar (N=21) rats were implanted with sterile radiotelemetry transmitters (Data Sciences International, St Paul, MN) in the abdominal cavity as previously described (Taffe et al., 2015; Wright et al., 2012). Animals were evaluated in clean standard plastic homecages (thin layer of bedding) in a dark testing room, separate from the vivarium, during the (vivarium) dark cycle. Radiotelemetry transmissions were collected via telemetry receiver plates (Data Sciences International, St Paul, MN; RPC-1 or RMC-1) placed under the cages as described in prior investigations (Aarde et al., 2013; Miller et al., 2013). Test sessions for inhalation studies started with a 15-minute interval to ensure data collection, then a 15-minute interval for baseline temperature and activity values followed by initiation of vapor sessions. The 15-minute baseline was omitted for the studies with intraperitoneal injection of THC due to the delayed onset of hypothermia (Nguyen et al., 2016b; Taffe et al., 2015). Drugs or vehicle for the antagonist studies were injected prior to, or after, THC administration as specified in the following experimental descriptions.

### 2.2. Intravenous catheterization

Groups of rats were anesthetized with an isoflurane/oxygen vapor mixture (isoflurane 5% induction, 1-3% maintenance) and prepared with chronic indwelling intravenous catheters as described previously (Aarde, Huang & Taffe, 2017; Miller, Aarde, Moreno, Creehan, Janda & Taffe, 2015; Nguyen, Grant, Creehan, Vandewater & Taffe, 2017). Briefly, the intravenous catheters consisted of a 14.5-cm length of polyurethane based tubing (Micro-Renathane®, Braintree Scientific, Inc, Braintree, MA) fitted to a guide cannula (Plastics One, Roanoke, VA) curved at an angle and encased in dental cement anchored to an ∼3 cm circle of durable mesh. Catheter tubing was passed subcutaneously from the animal’s back to the right jugular vein. Catheter tubing was inserted into the vein and tied gently with suture thread. A liquid tissue adhesive was used to close the incisions (3M™ Vetbond™ Tissue Adhesive: 1469SB, 3M, St. Paul, MN). A minimum of 4 days was allowed for surgical recovery prior to starting an experiment. For the first three days of the recovery period, an antibiotic (cefazolin, i.v.) and an analgesic (flunixin, s.c.) were administered daily. During testing, intravenous catheters were flushed with ∼0.2-0.3 ml heparinized (166.7 USP/ml) saline 2-3 times per week. Animals were excluded if catheter patency failure occurred (and no blood could be drawn) or if the external port had been chewed off by a cage mate.

### 2.3. Materials

Δ^9^-tetrahydrocannabinol (THC) was administered by vapor inhalation with doses described by the concentration in the propylene glycol (PG) vehicle and duration of inhalation as in prior studies (Javadi-Paydar et al., 2018; Nguyen et al., 2016b). THC was also administered intraperitoneally in a dose of 10 mg/kg, which produces robust temperature responses (Nguyen et al., 2016b; Taffe et al., 2015). SR141716 (Rimonabant; SR), AM-251 or AM-630 were administered intraperitoneally in a dose of 4 mg/kg. For injection, THC, SR, AM-251 or AM-630 were suspended in a vehicle of 95% ethanol, Cremophor EL and saline in a 1:1:8 ratio. The THC was provided by the U.S. National Institute on Drug Abuse; SR141716 was obtained from ApexBio (New Delhi, Delhi, India; Distributor: Fisher Scientific, Pittsburgh, PA, USA); AM-251 and AM-630 were obtained from Tocris Bioscience (Avonmouth, Bristol, UK; Distributor: Fisher Scientific, Pittsburgh, PA, USA).

### 2.4. Inhalation Apparatus

Sealed exposure chambers were modified from the 259 mm X 234 mm X 209 mm Allentown, Inc (Allentown, NJ) rat cage to regulate airflow and the delivery of vaporized drug to rats as has been previously described (Nguyen et al, 2016a; Nguyen et al, 2016b). An e-vape controller (Model SSV-1; La Jolla Alcohol Research, Inc, La Jolla, CA, USA) was triggered to deliver the scheduled series of puffs (4 10 second puffs, 2 second intervals, every 5 minutes) from Protank 3 Atomizer (Kanger Tech; Shenzhen Kanger Technology Co.,LTD; Fuyong Town, Shenzhen, China) e-cigarette cartridges by MedPC IV software (Med Associates, St. Albans, VT USA). The chamber air was vacuum-controlled by a chamber exhaust valve (i.e., a “pull” system) to flow room ambient air through an intake valve at ∼1 L per minute. This also functioned to ensure that vapor entered the chamber on each device triggering event. The vapor stream was integrated with the ambient air stream once triggered.

### 2.5. Nociception Assay

Tail-withdrawal antinociception was assessed using a water bath (Bransonic® CPXH Ultrasonic Baths, Danbury, CT) maintained at 52°C as previously described (Javadi-Paydar et al., 2018; Nguyen et al., 2016b). The latency to withdraw the tail was measured using a stopwatch, and a cutoff of 15 seconds was used to avoid any possible tissue damage (Wakley and Craft, 2011; Wakley et al, 2014). Tail-withdrawal was assessed 35, 60 and 120 minutes after the initiation of inhalation. The investigator performing the assay was blinded to the treatment condition for a given subject.

### 2.6. Plasma THC analysis

Blood samples were collected (∼300-500 µl) via jugular needle insertion following anesthesia with an isoflurane/oxygen vapor mixture (isoflurane 5% induction) or via implanted catheter for groups prepared for i.v. sampling. Plasma THC content was quantified using fast liquid chromatography/mass spectrometry (LC/MS) adapted from (Irimia, Polis, Stouffer, & Parsons, 2015; Lacroix & Saussereau, 2012; Nguyen et al., 2018). 50 µl of plasma were mixed with 50 µl of deuterated internal standard (100 ng/ml CBD-d3 and THC-d3; Cerilliant), and cannabinoids were extracted into 300 µL acetonitrile and 600 µl of chloroform and then dried. Samples were reconstituted in 100 µl of an acetonitrile/methanol/water (2:1:1). Separation was performed on an Agilent LC1100 using an Eclipse XDB-C18 column (3.5um, 2.1mm x 100mm) using gradient elution with water and methanol, both with 0.2 % formic acid (300 µl/min; 73-90%). Cannabinoids were quantified using an Agilent MSD6140 single quadrodpole using electrospray ionization and selected ion monitoring [CBD (m/z=315.2), CBD-d3 (m/z=318.2), THC (m/z=315.2) and THC-d3 (m/z=318.2)]. Calibration curves were conducted daily for each assay at a concentration range of 0-200 ng/mL and observed correlation coefficients were 0.999.

### 2.7. Experiments

#### 2.7.1. Experiment 1 (Nociception)

This study was conducted in groups of male (N=8) and female (N=7) Wistar rats that had previously been exposed to THC vapor inhalation (12.5-100 mg/mL; 30 minutes) to determine plasma levels of THC with dosing no more frequently than a 2 week interval necessary for recovery of blood volume after repeated sampling (Diehl et al., 2001). These groups were 19 weeks of age at the start of the present study. The female group was originally N=8 but one animal was lost to the study during surgical implantation of an intravenous catheter. For the present study, tail-withdrawal was assessed 35, 60 and 120 minutes after initiation of vapor inhalation of PG or THC (100 mg/mL for 30 minutes with 4 puffs every 5 minutes) with SR141716 (0, 4 mg/kg SR, i.p.) administered 15 minutes prior to PG or THC. Treatment order of these three conditions was counter-balanced within both male and female groups, with the vapor inhalations conditions conducted in same-sex pairs.

#### 2.7.2. Experiment 2 (SR141716 in male Sprague-Dawley and Wistar rats)

This study was conducted in a group of male Sprague-Dawley rats implanted with radiotelemetric devices used in the studies reported in a prior report on the efficacy of the vapor inhalation model (Nguyen et al., 2016b). This study was conducted immediately after Experiment 1 in that prior report (which evaluated the effect of vapor inhalation of THC (200 mg/mL) for 10, 20 or 30 minutes in otherwise naïve animals). In this study, animals received either the vehicle or SR141716 (4 mg/kg, i.p.) 15 minutes before the initiation of THC (200 mg/mL) inhalation for 20 minutes. This group next participated in a series of studies in which they were exposed to PG or THC (200 mg/mL) vapor for 20 or 30 minutes following injection of either the vehicle or SR141716 (4 mg/kg, i.p.). The pre-treatment interval was 90 minutes before the initiation of inhalation. These eight conditions were counter-balanced across the group with THC administered no more frequently than once per week. The group next participated in a study in which they were exposed to THC (200 mg/mL) vapor for 30 minutes and injected with either the vehicle or SR141716 (4 mg/kg, i.p.) after the inhalation session and prior to the start of telemetry recording. A final study was conducted in a group (N=13) of male Wistar rats used in previously reported stimulant vapor inhalation studies (Nguyen et al., 2016a). These rats received injection of either the vehicle or SR141716 (10 mg/kg, i.p.) 20 minutes before the initiation of THC (200 mg/mL) inhalation for 20 minutes.

#### 2.7.3. Experiment 3 (SR141716 after THC, i.p., in male Sprague-Dawley rats)

This study was conducted to determine if SR141716 could reverse hypothermia induced by THC injection, using two groups (N=8 per group) of male Sprague-Dawley rats. SR141716 (4 mg/kg, i.p.), or vehicle, was injected 45 minutes after the injection of THC (20 mg/kg, i.p.) in the group used in Experiment 2. A study of the effect of SR141716 (4 mg/kg, i.p.), or vehicle, injected 90 minutes after THC (20 mg/kg, i.p.), or vehicle, was conducted in a different group of male Sprague-Dawley rats that was received in the laboratory and implanted with telemetry transmitters at the same time as the first group, but used in studies of the effect of psychomotor stimulants 3,4-methylenedioxymethpyrovalerone and alpha-pyrrolidinopentiophenone, by i.p. injection (unpublished data).

#### 2.7.4. Experiment 4 (THC, i.v., in male and female Wistar rats)

This study was conducted in groups of naive male (N=8) and female (N=7) Wistar rats implanted with radiotelemetric devices and intravenous catheters. The goal was first to determine the timecourse of any thermoregulatory or activity effects, compared with the profile following inhaled or i.p. injected THC which differ significantly in our ongoing studies. A rapid decline of temperature to a nadir is observed within about 60 minutes of initiating vapor inhalation and a recovery within about 3 hours is contrasted with a slow decline to a nadir approximately 5-6 hours after i.p. injection. Treatments of THC (0.0, 0.1, 0.3 and 1.0, i.v.) were therefore evaluated in a counterbalanced order on two occasions, once with and once without serial blood sampling, with THC exposure no more frequently than once per week. Blood sampling experiments were conducted no more frequently then every other week to accord with recommendations for blood volume recovery (Diehl et al., 2001) and rats were injected with ∼300-500 µl of saline, i.v., after each blood sample was obtained to maintain circulating volume. Only the experiments in which THC or vehicle was administered without any blood sampling are included in this report. Telemetry function was lost in 2 female rats, thus N=5 for the analysis. Following these initial studies, the groups completed a study in which they received THC (0.0, 1.0 mg/kg, i.v.) with SR141716 (0.0, 4.0 mg/kg, i.p.) for four total treatment conditions in a counterbalanced order.

#### 2.7.5. Experiment 5 (Antagonist Challenge After Repeated THC Vapor Inhalation)

Human laboratory studies of cannabinoid effects are often conducted with cannabis-experienced individuals who may be partially tolerant to the effects of THC. We’ve recently shown that repeated-THC vapor exposure produces immediate tolerance in female rats (Nguyen et al., 2020; Nguyen et al., 2018) that lasts into adulthood after adolescent treatment (Nguyen et al., 2020). In this experiment a twice-daily repeated-THC exposure study was conducted with a group of female rats (N=8; 11 weeks of age at start of this study) implanted with radiotelemetric devices. This provided the ability to monitor temperature responses without the handling stress of repeated rectal assessment and facilitated a more precisely determined timecourse of tolerance as it developes across the 7 sessions in adult female rats (Nguyen et al., 2018). Inhalation sessions were 30 minutes in duration and the THC concentration was 100 mg/ml in this study. In the first week, rats received four days of twice-daily inhalation of the PG vehicle (5.25 h interval between session initiations on each day). The following week, rats received five days of twice daily inhalation of THC (100 mg/mL), again with 5.25 h between sessions. Additional THC-inhalation sessions were conducted one and two weeks after the final session of the repeated-THC week to determine if tolerance persisted.

The next study was conducted starting 20 days after the final post-chronic THC-inhalation assessment to determine if the hypothermic effects of THC inhalation were mediated by CB_1_ and/or CB_2_ receptors using pretreatment with 4 mg/kg, i.p. of the CB_1_ antagonists/inverse agonists SR141716 and AM-251 and the CB_2_ antagonist/inverse agonist AM-630. In these studies, the THC exposures were no more frequent than a 7 day interval and the order of the pre-treatment / inhalation conditions was counter-balanced across the group within each antagonist study. The final study was conducted in the same group, to determine the effect of SR141716 (4 mg/kg, i.p.) and AM-251 (4 mg/kg, i.p.) on the hypothermic effect of injected THC (10 mg/kg, i.p.). For these studies, the vehicle or one of the two antagonists were administered 15 minutes prior to injection with THC or Vehicle, with treatment conditions evaluated in a counterbalanced order.

#### 2.7.6. Experiment 6 (Plasma THC by sex and strain of rat)

Although we’ve previously reported plasma THC levels after vapor inhalation for male and female Wistar rats (Javadi-Paydar et al., 2018; Nguyen et al., 2018), and for Wistar and Sprague-Dawley male rats (Taffe et al., 2020), the results are not directly cross-comparable and in most cases were for single time-points after inhalation. Therefore it was of interest in this study to directly compare plasma THC across the post-administration interval over which body temperature is affected, using age-matched groups. To this end, male and female Wistar and Sprague-Dawley rats (N=6 per group) were prepared with intravenous catheters and permitted to recover for at least a week. Blood samples (∼300-500 µl) were obtained through the catheters 35, 60, 120, and 240 minutes after the start of inhalation of THC (100 mg/mL) for 30 minutes. In the next experiment, conducted 5 weeks later, blood samples were obtained at the same time points following injection of THC (10 mg/kg, i.p.); catheters remained patent in only 3 female rats of each strain in the i.p. study thus the female data are not reported. Rats were injected with ∼300-500 µl of saline, i.v., after each blood sample was obtained to maintain circulating volume. An experiment involving blood sampling on a similar schedule after a 30 minute session of nicotine (30 mg/mL) vapor inhalation was interposed between these two THC experiments. Blood sampling experiments were conducted no more frequently then every other week to accord with recommendations for blood volume recovery (Diehl et al., 2001).

### 2.8. Data Analysis

The telemeterized body temperature and activity rate (counts per minute) were collected on a 5-minute schedule in telemetry studies but are expressed as 30 minute averages for analysis. The time courses for data collection are expressed relative to the THC injection time or the initiation of vapor inhalation and times in the figures refer to the end of the interval (e.g. “60 minutes” reflects the data collected from 35 to 60 minutes, inclusive). Any missing temperature values were interpolated from the values before and after the lost time point. Activity rate values were not interpolated because 5-minute to 5-minute values can change dramatically, thus there is no justification for interpolating. Group sizes were determined initially with a power analysis of similar data collected previously in the laboratory. Data (tail-withdrawal latency, telemeterized temperature and activity measures, plasma THC concentrations) were generally analyzed with two-way Analysis of Variance (ANOVA) including repeated measures factors for the Drug treatment condition and the Time after vapor initiation or injection. A mixed-effects model was used instead of ANOVA wherever there were missing datapoints. Any significant main effects were followed with post-hoc analysis using Tukey (multi-level factors) or Sidak (two-level factors) procedures. All analysis used Prism 8 for Windows (v. 8.4.2; GraphPad Software, Inc, San Diego CA).

## 3. Results

### 3.1. Experiment 1 (Nociception)

This experiment confirmed that the CB1 antagonist SR141716 partially (female rats) or completely (male rats) blocked the anti-nociceptive effect of THC inhalation. The analysis of the female rat data (**Figure 1**) confirmed a significant effect of Drug treatment condition [F (2, 12) = 23.01; P<0.0001] but not of Time after vapor initiation or of the interaction of factors. Post-hoc analysis of the marginal means confirmed that tail-withdrawal latency differed significantly across all three treatment conditions. The analysis of the male rat data confirmed a significant effect of Drug treatment condition [F (2, 14) = 13.54; P=0.0005], of Time after vapor initiation [F (2, 14) = 5.85; P=0.0142] but not of the interaction of factors. The Tukey post-hoc analysis confirmed that tail-withdrawal latency was significantly longer after the Veh-THC condition compared with the SR-PG (35-60 minutes after initiation of inhalation) and SR-THC (35 minutes post-initiation) conditions.

**Figure 1:**
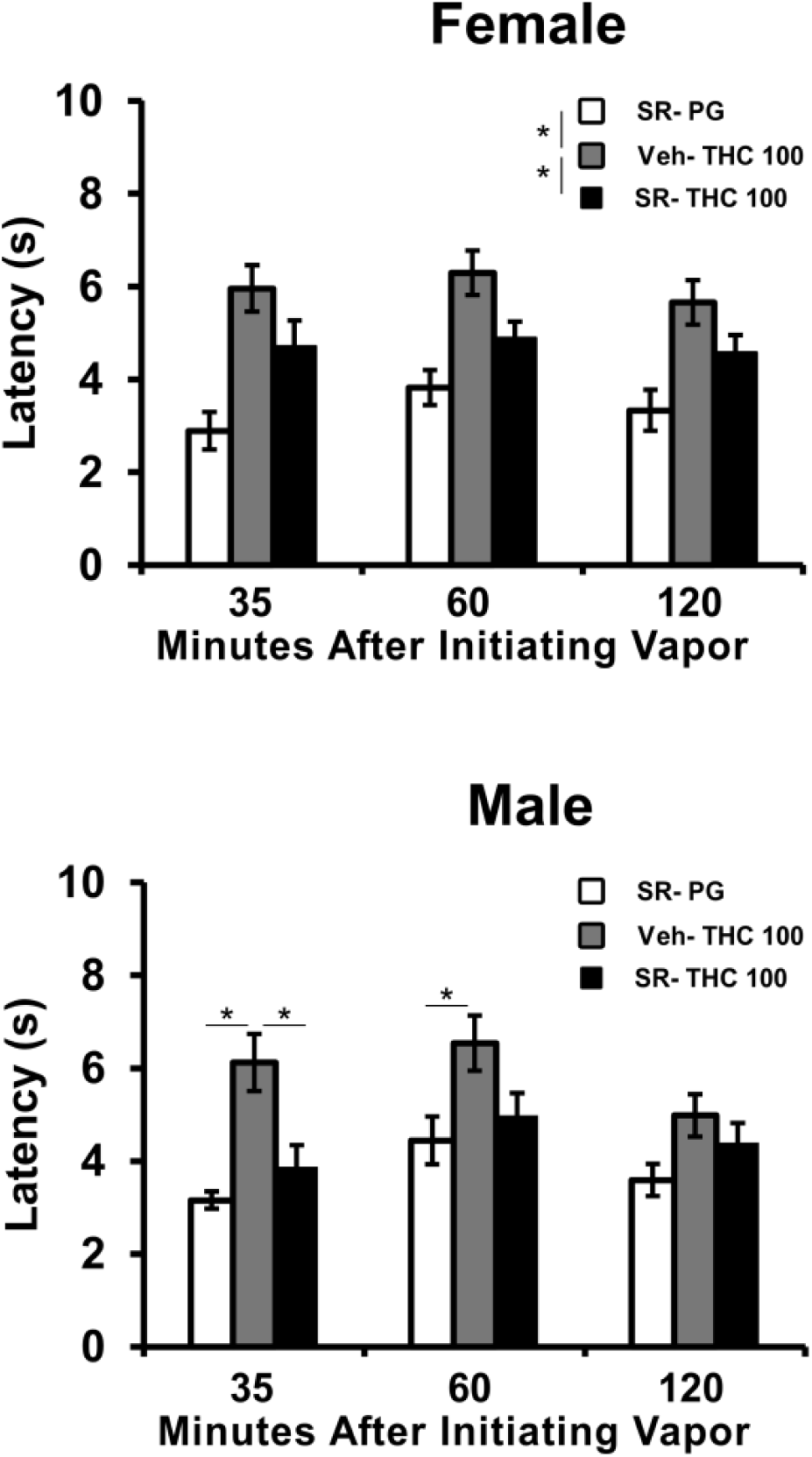
Mean tail-withdrawal latencies for female (N=7; ±SEM) and male (N=8; ±SEM) rats after pre-treatment with SR141716 (SR; 4 mg/kg, i.p.) or the vehicle (Veh) and inhalation of PG or THC (100 mg/mL) vapor. Significant differences between pairs of conditions or the marginal means are indicated with *.

### 3.2. Experiment 2 (SR141716 in male Sprague-Dawley and Wistar rats)

Treatment of male Sprague-Dawley rats (N=8) with 4 mg/kg SR141716, i.p.,15 minutes prior to THC (200 mg/L) inhalation for 20 minutes slightly altered the course of the hypothermic response (**Figure 2A**). The ANOVA confirmed significant effects of Time post-initiation [F (6, 42) = 29.65; P<0.0001] and of the interaction of Time post-initiation with the pre-treatment condition [F (6, 42) = 3.57; P<0.01]. The Tukey post-hoc test confirmed that temperature was significantly lower than both pre-inhalation time points for Veh + THC (30-90 minutes after vapor initiation) and for SR141716 + THC (30-60 minutes after vapor initiation). The post-hoc test further confirmed that body temperature was higher 120 minutes after vapor initiation when SR141716 was administered compared with the vehicle.

**Figure 2:**
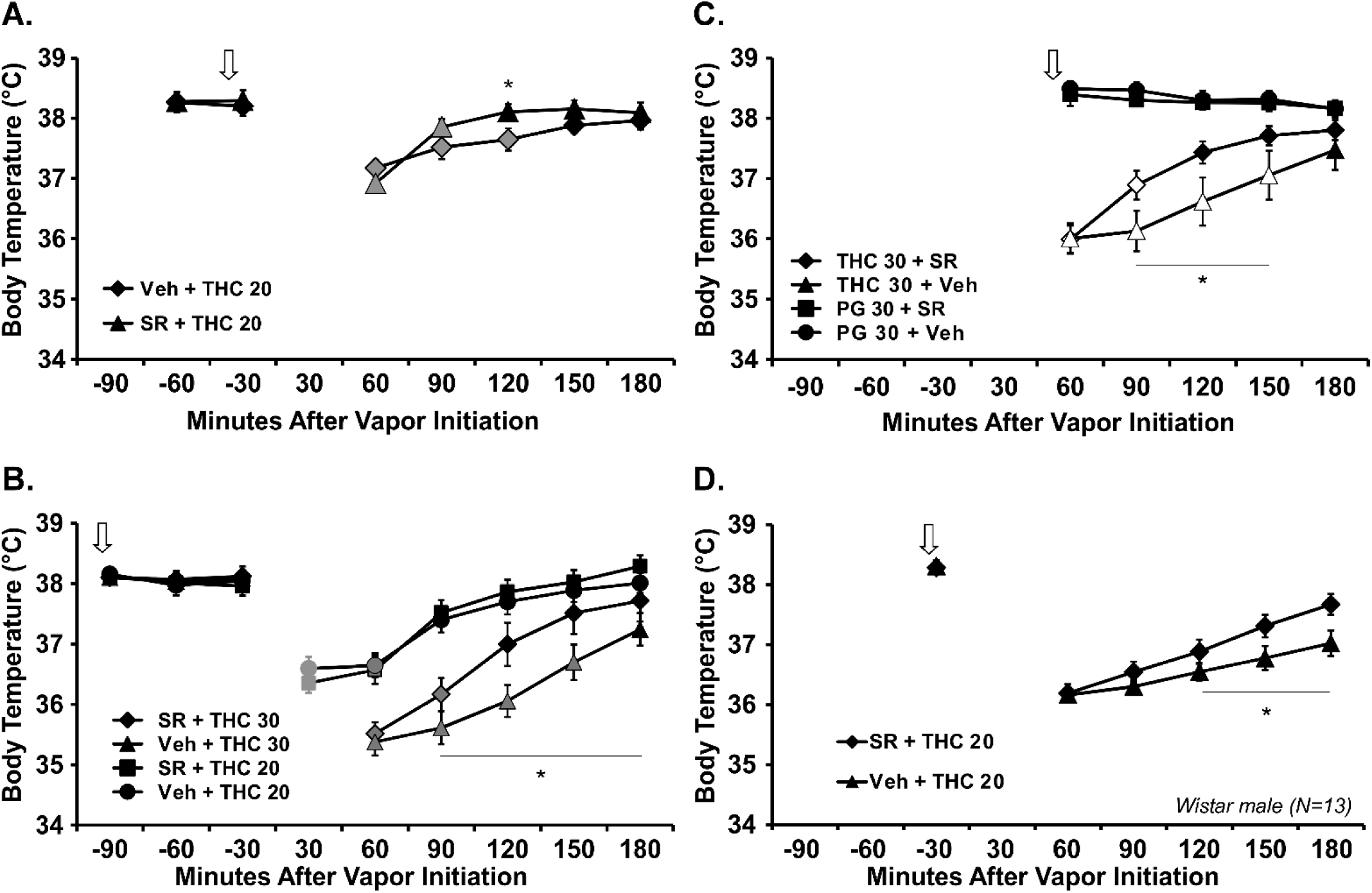
Mean body temperature of male A-C) Sprague-Dawley (N=8; ±SEM) and D) Wistar (N=13; ±SEM) rats following vapor inhalation of THC (200 mg/mL) for 20 (THC 20) or 30 (THC 30) minutes. The effect of the CB1 antagonist SR141716 (SR; 4 mg/kg, i.p.) was assessed for the Sprague-Dawley rats A) 15 minutes before vapor initiation; B) 90 minutes prior to vapor initiation and C) after vapor cessation. D) The effect of (SR; 10 mg/kg, i.p.; 20 minutes prior to inhalation) on temperature disruption associated with THC (200 mg/mL) inhalation for 20 minutes was assessed for the Wistar rats. Arrows indicate the timing of the vehicle/SR141716 injection. Shaded symbols indicate a significant difference from the baseline value, and a significant difference between SR141716 (SR) pre-treatment conditions is indicated with *. The PG vapor inhalation condition was used in lieu of a baseline measure for confirmation of hypothermia in the post-vapor SR141716 experiment, thus open symbols indicate a significant difference from the respective PG inhlation condition.

Increasing the pre-treatment interval so that SR141716 was administered 90 min prior to the initiation of vapor inhalation (THC 200 mg/mL; 20 vs 30 minutes) resulted in a profile of effect (**Figure 2B**) that was similar to that produced by the 15 min pretreatment interval. The three-way ANOVA confirmed significant effects of Inhalation Duration [F (1, 168) = 88.42; P<0.0001], of Time Post-initiation [F (5, 168) = 41.58; P<0.0001] and of Vehicle / SR pre-treatment condition [F (1, 168) = 9.18; P=0.0028]. The 30 minute timepoint for the shorter inhalation condition was omitted in this analysis because it did not exist for the longer inhalation condition. There was also a significant effect of the interaction of Inhalation Duration with Time post-initiation [F (5, 168) = 6.14; P<0.0001] and the interaction of Inhalation Duration with Vehicle / SR141716 pre-treatment condition [F (1, 168) = 4.42; P<0.05] on the body temperature. To further explore the interaction, a two-way ANOVA on the 30 minute inhalation conditions confirmed significant effects of Vehicle / SR141716 pre-treatment [F (1, 7) = 6.13; P=0.0425], Time post-initation [F (5, 35) = 57.20; P<0.0001] and the interaction [F (5, 35) = 6.56; P=0.0002]. The post-hoc test confirmed a significant effect of SR141716 on body temperature from 90-180 minutes after vapor initiation. The temperature data for the PG inhalation conditions and the activity data for the whole study are shown in **Supplemental Figure S1**.

Administration of SR141716 (4 mg/kg, i.p.) at the end of a 30 minute THC (200 mg/mL) vapor inhalation session likewise modified the course of hypothermia (**Figure 2C**). The three-factor ANOVA confirmed significant effects of Time after vapor initiation [F (4, 140) = 6.37; P<0.0001], of Vapor condition [F (1, 140) = 197.8; P<0.0001], of Vehicle / SR141716 Post-treatment condition [F (1, 140) = 4.87; P<0.05], of the interaction of Time with Vapor condition [F (4, 140) = 11.86; P<0.0001], and the interaction of Vapor condition with Post-treatment condition [F (1, 140) = 8.53; P<0.005] on body temperature. To further parse the effect of SR141716 post-treatment a followup ANOVA was conducted on only the THC-inhalation conditions and the ANOVA confirmed significant effects of Time after vapor initiation [F (5, 35) = 41.97; P<0.0001], and of the interaction of Time with Post-treatment condition [F (5, 35) = 5.30; P=0.001], on body temperature. The posthoc test confirmed that temperature differed between post-treatment condition from 90-150 minutes after the start of inhalation.

Increasing the SR141716 dose to 10 mg/kg, i.p., administered 15 min prior to the start of inhalation (THC 200 mg/mL; 20 minutes) in male Wistar rats (**Figure 2D**) did not produce a qualitatively different outcome. The analysis confirmed a significant effect of Time Post-initiation [F (5, 60) = 61.23; P<0.0001] and of interaction of Time Post-initiation with Vehicle / SR141716 pre-treatment condition [F (5,60) = 6.60; P<0.0001] on body temperature. The posthoc test confirmed that temperature differed between pre-treatment conditions from 120-150 minutes after the start of inhalation.

### 3.3. Experiment 3 (SR141716 after THC, i.p., in male Sprague-Dawley rats)

This experiment confirmed that a 4 mg/kg, i.p. SR141716 dose was capable of affecting the hypothermic response to THC injection when administered 45 minutes (**Figure 3A**) or 90 minutes (**Figure 3B**) after the initiation of vapor inhalation, attenuating the response when the body temperature response was first initiated (45 minutes) and reversing it when fully engaged (90 minutes). The ANOVAs confirmed significant effects of Drug Condition, Time post-injection and the interaction on body temperature for the 45 minute [Drug: F (1, 7) = 12.57; P<0.01; Time: F (9, 63) = 21.8; P<0.0001; P<0.0001; Interaction: F (9, 63) = 21.77; P<0.0001] and 90 minute [Drug: F (3, 21) = 45.29; P<0.0001; Tim e: F (9, 63) = 22.65; P<0.0001; P<0.0001; Interaction: F (27, 189) = 25.27; P<0.0001] studies. The SR141716 did not affect activity rate when administered 45 minutes after THC (**Figure 3C**), as there was no significant effect of Drug condition or interaction with Time post-injection, there was only a main effect of Time [F (9, 63) = 16.84; P<0.0001] on activity rate. In the case of the 90 minute study, the SR reversed the locomotor suppression caused by THC injection (**Figure 3D**). The ANOVA confirmed significant effects of Drug Condition [F (3, 21) = 28.72; P<0.0001], Time post-injection [F (9, 63) = 11.88; P<0.0001] and the interaction [F (27, 189) = 2.43; P=0.0003] on activity rate. Thus, SR141716 was proved efficacious at *reversing* the effects of THC when injected, using identical methods to record temperature and activity as used in the vapor inhalation study.

**Figure 3:**
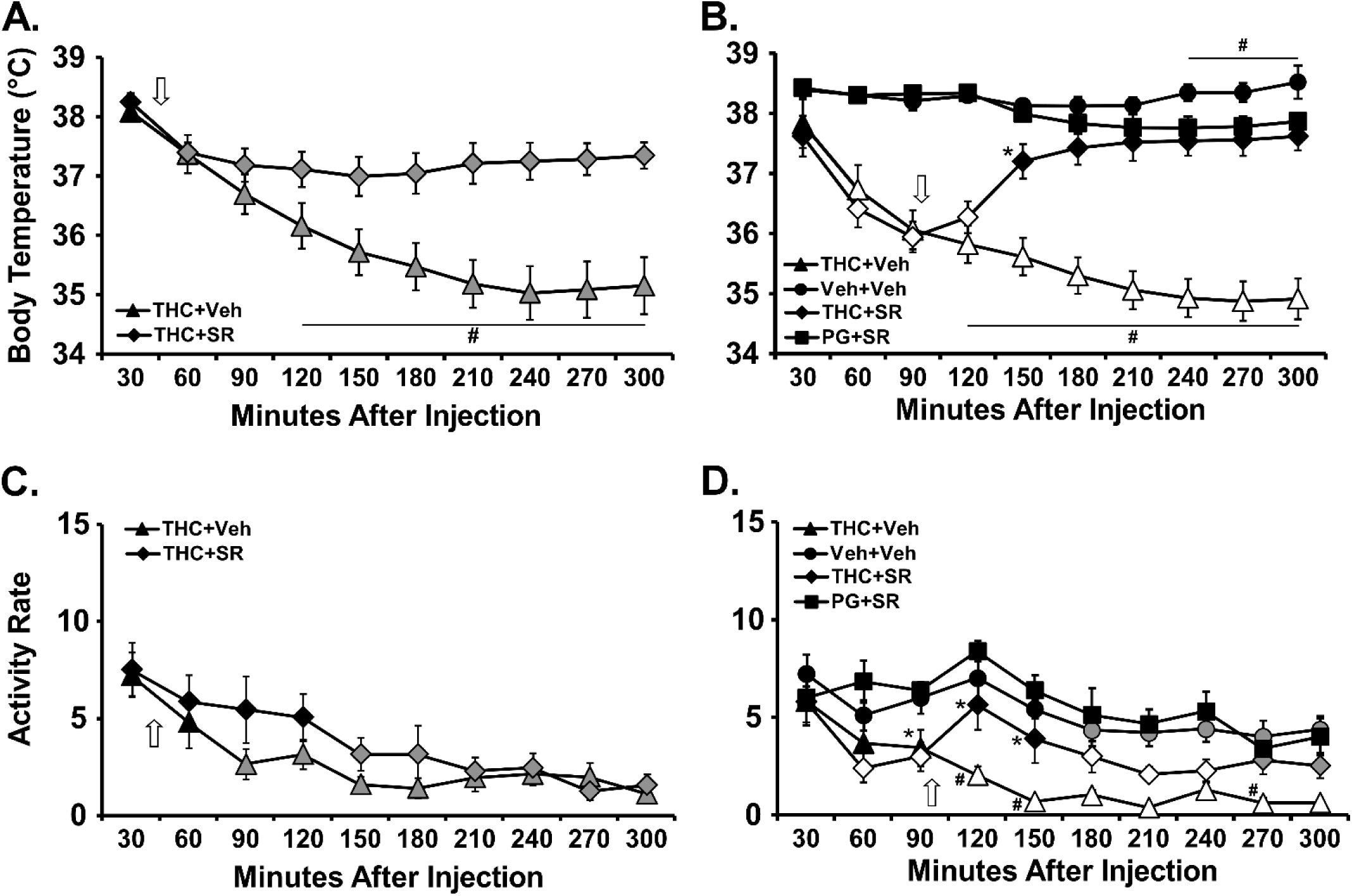
Mean (N=8; ±SEM) temperature (A, B) and Activity rate (C, D) for groups of rats injected (i.p.) with THC (20 mg/kg) and then SR141716 (4 mg/kg) or the vehicle (Veh) either 45 (A, C) or 90 (B, D) minutes later. Arrows indicate the timing of the vehicle/SR141716 injection. The vehicle control for THC was included in the 90 minute study only. Shaded symbols indicate a significant difference from the 30 minute value, within treatment condition. Open symbols indicate a significant difference from the 30 minute value and the respective Vehicle condition; a difference from the Vehicle only is indicated with *. A significant difference between SR141716 (SR) treatment conditions is indicated with #.

### 3.4. Experiment 4 (THC, i.v., in male and female Wistar rats)

Temperature was decreased rapidly after i.v. injection in both female (1.0 mg/kg, i.v.) and male (0.3, 1.0 mg/kg, i.v.) Wistar rats (**Figure 4A**,**B**). One male rat’s catheter became non-patent after completing only the vehicle and THC 0.1 mg/kg doses. One female rat’s transmitter never functioned properly. Catheter failure in two of the remaining individuals meant that four female rats contributed to each dose condition and in aggregate it was N=6 for all conditions except N=5 for the 1.0 mg/kg dose condition. The mixed-effects analysis confirmed significant effects of Time [F (8, 48) = 11.07; P<0.0001] and Dose [F (3, 18) = 9.607; P=0.0005] for female rats and of Time [F (8, 56) = 24.54; P<0.0001], Dose [F (3, 21) = 14.65; P<0.0001] and the interaction [F (24, 150) = 3.48; P<0.0001] for the male rats, on body temperature. Activity was reduced in the female group after the 1.0 mg/kg, i.v., dose but the mixed-effects analysis failed to confirm any significant effects of treatment condition on activity in female [Time post-injection: F (8, 48) = 9.06; P<0.0001] or male [Time post-injection: F (8, 56) = 19.91; P<0.0001] groups (**Figure 4C,D**). The rats had become somewhat tolerant to the effect of 1.0 mg/kg, i.v. by the time of the SR141716 experiment but significant reductions in body temperature were still observed in the vehicle pretreatment, 1.0 mg/kg THC, i.v., condition for both male and female groups (**Figure 4E,F**). Seven male rats and four female rats had both functional transmitters and catheters for the entire SR141716 study. The ANOVAs confirmed that SR141716 pre-treatment attenuated the effects of THC on body temperature in female [Time: F (8, 24) = 17.88; P<0.0001; Dose Condition: F (3, 9) = 9.32; P<0.005; Interaction: F (24, 72) = 2.99; P<0.0005] and male [Time: F (8, 48) = 19.00; P<0.0001; Dose Condition: F (3, 18) = 3.85; P<0.05] rats. There were no effects of THC on activity rate in either sex (not shown).

**Figure 4:**
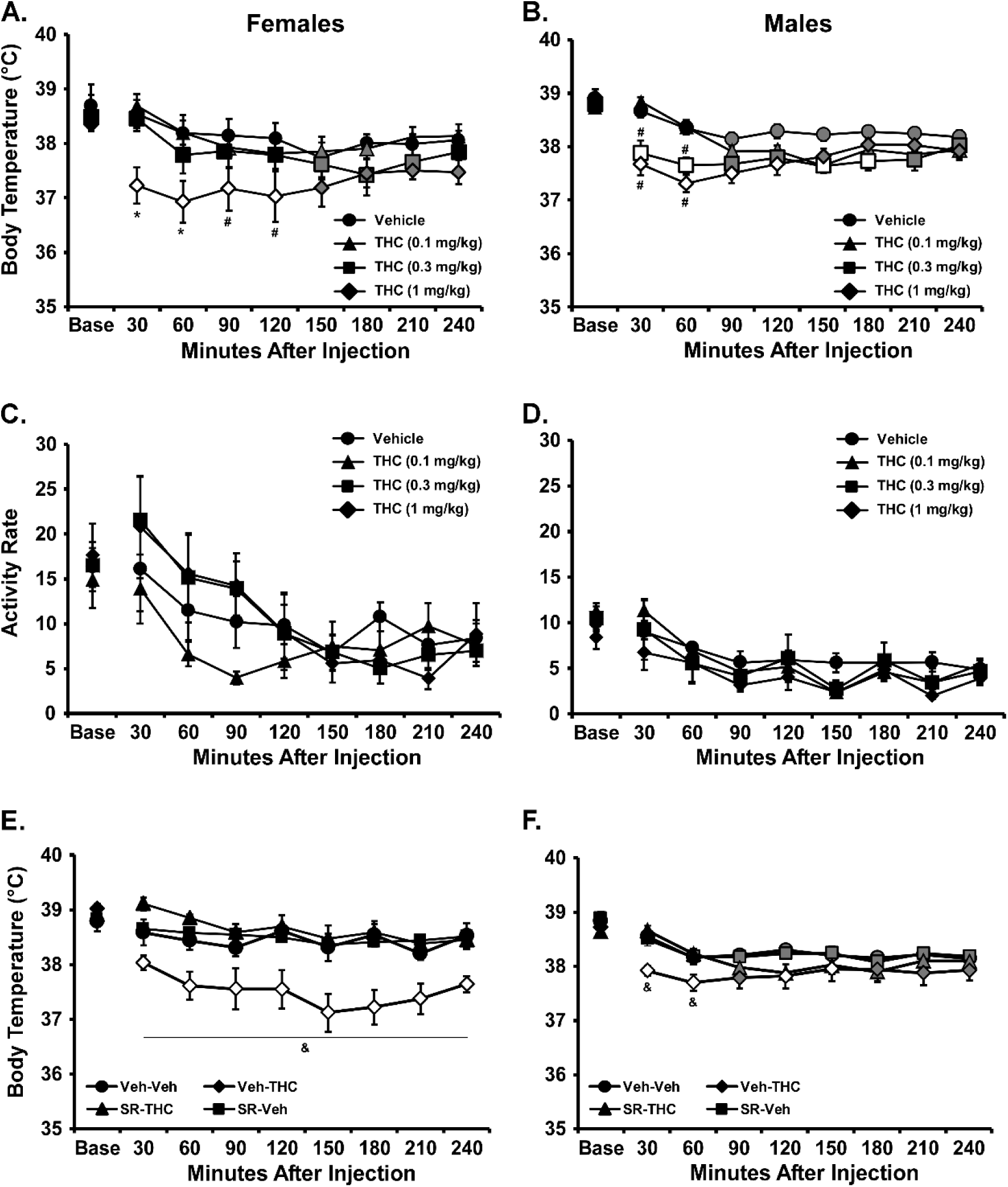
Mean (±SEM) temperature (A, B) and Activity (C, D) response to THC injection (i.v.) in female (A, C) and male (B, D) rats. Group sizes range from N=5-8, see text. Mean (±SEM) temperature response to THC injection (i.v.) in E) female (N=4) and F) male (N=7) rats after injection (i.p.) with SR141716 (SR) or the vehicle (Veh).Shaded symbols indicate a significant difference from the baseline (Base), within treatment condition. Open symbols indicate a significant difference from the baseline and the respective Vehicle (i.v.) condition at a given time after injection. A significant difference from all other treatment conditions at a given time after injection is indicated with * and a difference from the 0.1 mg/kg condition with #. A significant difference between pre-treatment conditions, within vapor inhalation conditions, is indicated with &.

### 3.5. Experiment 5 (Antagonist Challenge After Repeated THC Vapor Inhalation)

Naïve female rats first exposed to four days of twice-daily inhalation of PG (30 minutes) exhibited no acute reduction in body temperature post-initiation of vapor as is shown in **Figure 5A.** In contrast body temperature was acutely reduced by inhalation of THC (100 mg/mL; 30 minutes) during the repeated-THC week (**Figure 5B**). The analysis of variance was conducted on 30-minute bins and included the first session of the day for all nine treatment days. The ANOVA confirmed significant effects of Session [F (8, 56) = 17; P<0.0001], of Time after vapor initiation [F (9, 63) = 19.7; P<0.0001] and of the interaction of factors [F (72, 504) = 9.98; P<0.0001]. The Tukey post-hoc test (see **Figure 5** for details) confirmed that there was no reduction in body temperature relative to baseline in the first 180 minutes after the initiation of inhalation for any of the PG sessions. In contrast the inhalation of THC induced a consistent hypothermia relative to the pre-inhalation baseline up to at least 210 minutes after vapor initiation for all five days. Similarly, the post-hoc confirmed that body temperature was lower compared with all four PG sessions 60 minutes after initiation of vapor in all five THC sessions. This continued for up to 180 minutes after initiation on the first two THC sessions. THC-associated hypothermia was significantly attenuated in initial magnitude on THC days 4 and 5 compared with the first three THC days. There were two additional THC sessions conducted 1 and 2 weeks after the repeated THC week (see **Supplemental Figure S2**).

**Figure 5:**
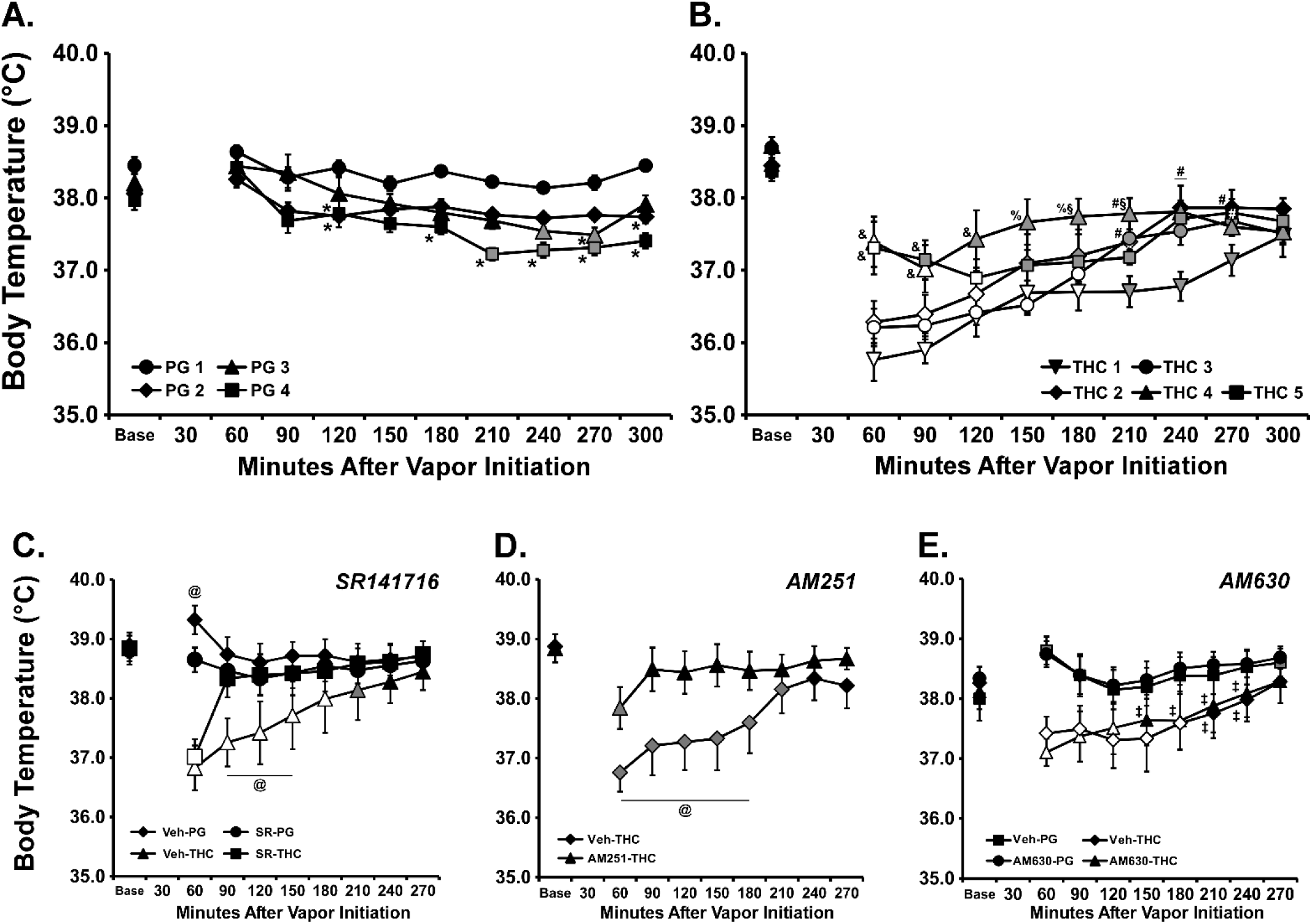
Mean (N=8; ±SEM) body temperature of female Wistar rats before (Base= baseline) and after vapor inhalation of A) PG or B) THC (100 mg/mL) for 30 minutes. Shaded symbols indicate a significant difference from the within-day baseline value, open symbols indicate a significant difference from the baseline and the respective time during **all four** PG inhalation days. A significant difference from PG day 1 is indicated by *, a difference from THC day 1 by #, a significant difference from THC days 1-3 by &, a significant difference from THC days 1 and 3 by %, and a significant difference between THC 4 and THC 5 by §; a bar indicates all other days differ. Mean (N=8; ±SEM) body temperature of female Wistar rats before (Base= baseline) and after vapor inhalation of PG or THC (100 mg/mL) for 30 minutes. Experiments included pretreatment of animals with CB1 antagonist/inverse agonists C) SR141716 and D) AM-251 as well as E) the CB2 antagonist AM-630. Shaded symbols indicate a significant difference from the baseline value, open symbols indicate a significant difference from the baseline and the respective time in the corresponding PG inhalation condition (when run). A significant difference from the respective PG condition only is indicated with ‡. A significant difference between vehicle and antagonist pre-treatment conditions is indicated with @.

Additional studies were conducted to probe the dependence of the THC-induced hypothermia on the CB_1_ or CB_2_ endocannabinoid receptor subtypes, starting 20 days after the prior THC session. For all these experiments the rats were exposed to THC no more frequently than a 7 day interval. In the first experiment (**Figure 5C**), SR141716 (4 mg/kg, i.p.) or the vehicle was administered 15 minutes prior to the start of vapor inhalation of PG or 100 mg/mL THC for 30 minutes. The four conditions were conducted in randomized order and the ANOVA confirmed a significant effect of Time after vapor initiation [F (8, 56) = 7.38; P<0.0001], of Treatment condition [F (3, 21) = 8.44; P<0.001] and of the interaction [F (24, 168) = 8.08; P<0.0001]. The Tukey post-hoc test confirmed a significant difference between Vehicle and SR141716 pretreatment conditions from 90-150 minutes after THC vapor initiation and 60 minutes after PG vapor initiation.

In the next experiment AM-251 (4 mg/kg, i.p.) or the vehicle was administered 15 minutes prior to start of vapor inhalation of 100 mg/mL THC for 30 minutes (**Figure 5D**). The two conditions were conducted in randomized order. The ANOVA confirmed a significant effect of Time after vapor initiation [F (8, 56) = 11.65; P<0.0001], of Treatment condition [F (1, 7) = 18.73; P<0.005] and of the interaction of factors [F (8, 56) = 6.38; P<0.0001]. The Tukey post-hoc test confirmed a significant difference between Vehicle and AM-251 pre-treatment for 60-180 minutes after the start of vapor. Finally, AM-630 (4 mg/kg, i.p.) or the vehicle was administered 15 minutes prior to start of vapor inhalation of PG or 100 mg/mL THC for 30 minutes (**Figure 5E**). The four conditions were conducted in randomized order and the ANOVA confirmed a significant effect of Time after vapor initiation [F (8, 56) = 4.59; P<0.0005], of Treatment condition [F (3, 21) = 23.62; P<0.0001] and of the interaction [F (24, 168) = 4.24; P<0.0001]. The Tukey post-hoc test confirmed that body temperature was significantly lower compared with both the baseline and the respective PG inhalation condition for Veh-THC (30-180 minutes after initation of vapor) and for AM-630-THC (30-120 minutes). Body temperature was significantly lower after THC inhalation compared the respective PG conditions up to 240 minutes after the start of inhalation.

For this study the female rats were injected with THC (0, 10 mg/kg i.p.) with the vehicle, SR141716 (4 mg/kg, i.p.) or AM-251 (4 mg/kg, i.p.) administered 15 minutes beforehand. The analysis of all six treatment conditions confirmed a significant effect of Time post-injection [F (9, 63) = 22.18; P<0.0001], of Drug treatment condition [F (5, 35) = 4.15; P<0.005] and of the interaction [F (45, 315) = 3.37; P<0.0001] on body temperature (**Figure 6**). The Tukey post-hoc test confirmed that temperature was lower than baseline and the respective time points after vehicle-vehicle injection when THC was preceded by the vehicle (30-270 minutes post-injection), when THC was preceded by AM-251 (60-270 minutes post-injection) and when the vehicle was preceded by AM-251 (90-120 minutes post-injection). Temperature was lower in the Veh-THC condition compared with the SR-THC treatment from 30-270 minutes after the THC injection. No significant differences in temperature were confirmed for the SR-Veh/SR-THC or AM-251-Veh/AM-251-THC pairs of conditions.

**Figure 6.**
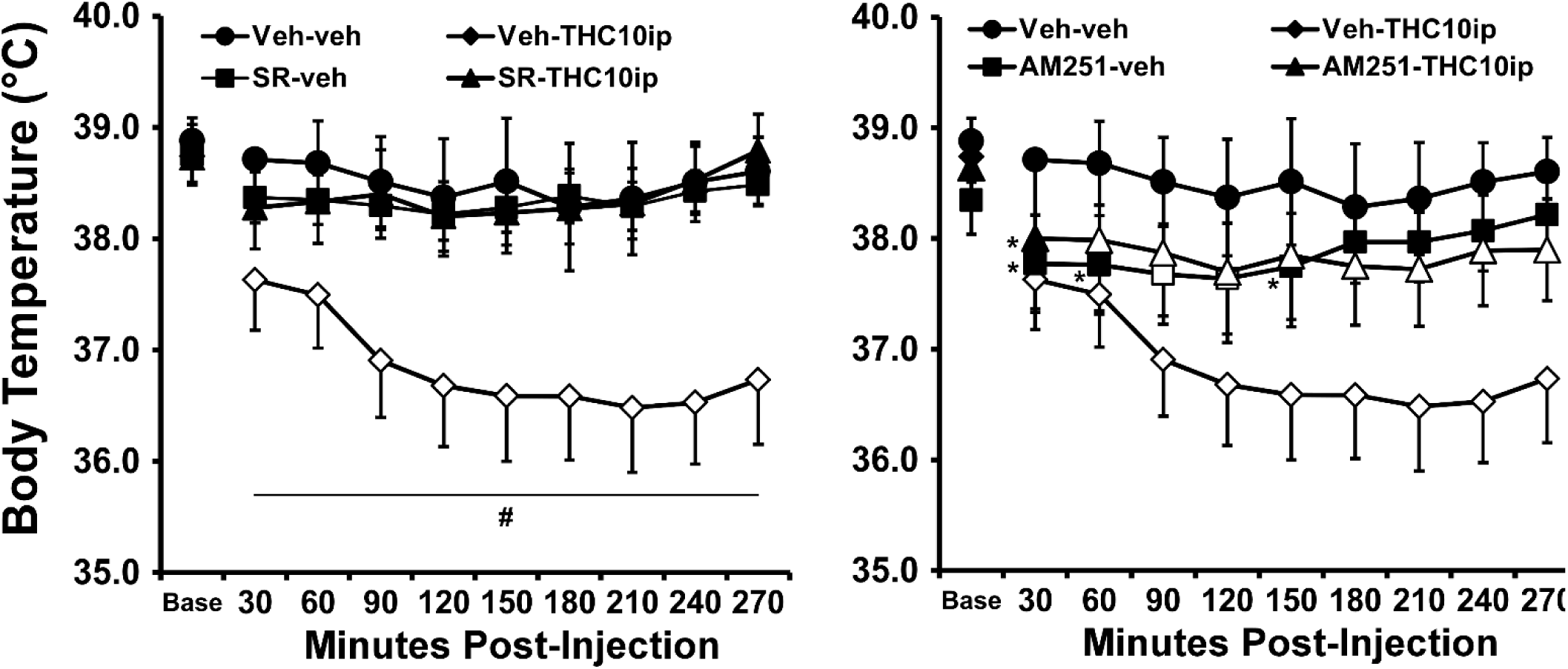
Mean (N=8; ±SEM) body temperature of female Wistar rats after i.p. injection of THC (10 mg/kg) or the Vehicle following pre-treatment with left) SR141716 (4 mg/kg, i.p.) or right) AM-251 (4 mg/kg, i.p.). The Vehicle-Vehicle and Vehicle-THC conditions are depicted in both panels to facilitate comparison. Shaded symbols indicate a significant difference from the baseline value, open symbols indicate a significant difference from the baseline and the Vehicle-Vehicle condition. A significant difference from the Vehicle-Vehicle condition, but not the baseline, is indicated with * and a difference between Veh and SR141716 pre-treatment THC conditions is indicated with #.

### 3.6. Experiment 6 (Plasma THC by sex and strain of rat)

The thermoregulatory studies presented cross rat age, strain, sex and prior treatment demonstrating how well the effects of CB1 antagonism generalize. Although we’ve shown previously that male and female Wistar rats achieve similar plasma levels of THC under identical inhalation conditions (Javadi-Paydar et al., 2018), as do male Wistar and Sprague-Dawley rats across ∼1 year of age (Taffe et al., 2019), this sex by strain comparison is necessarily indirect. The present experiment addressed that by treating age matched male and female rats of Wistar and Sprague-Dawley strains with the same inhalation conditions.

The plasma THC concentrations were analyzed first by Group and by Time after vapor initiation (**Figure 7**). The mixed-effects analysis confirmed a significant effect of Time [F (3, 51) = 55.13, P<0.0001], and of the interaction of Group with Time [F (9, 51) = 2.51, P<0.05]. The post-hoc test of the marginal means confirmed that plasma differed between all time points, save that plasma at 120 minutes did not differ from the levels at either 60 or 240 minutes after vapor initiation. The only Group differences confirmed at a specific time were between female WI and each male Group, 35 minutes after vapor initiation.To further explore the interaction, the data were analyzed first collapsed across strain and then collapsed across sex. The mixed-effects analyses confirmed a significant effect of Time [F (3, 57) = 54.86; P<0.0001) and the interaction of Sex with Time [F (3, 57) = 5.21; P<0.005] on plasma THC concentration when Strain was ignored and of Time [F (3, 57) = 45.86; P<0.0001], but not of Strain or of the interaction, when Sex was ignored. Again, the post-hoc test confirmed the sex difference only at the 35 minute time point.

**Figure 7:**
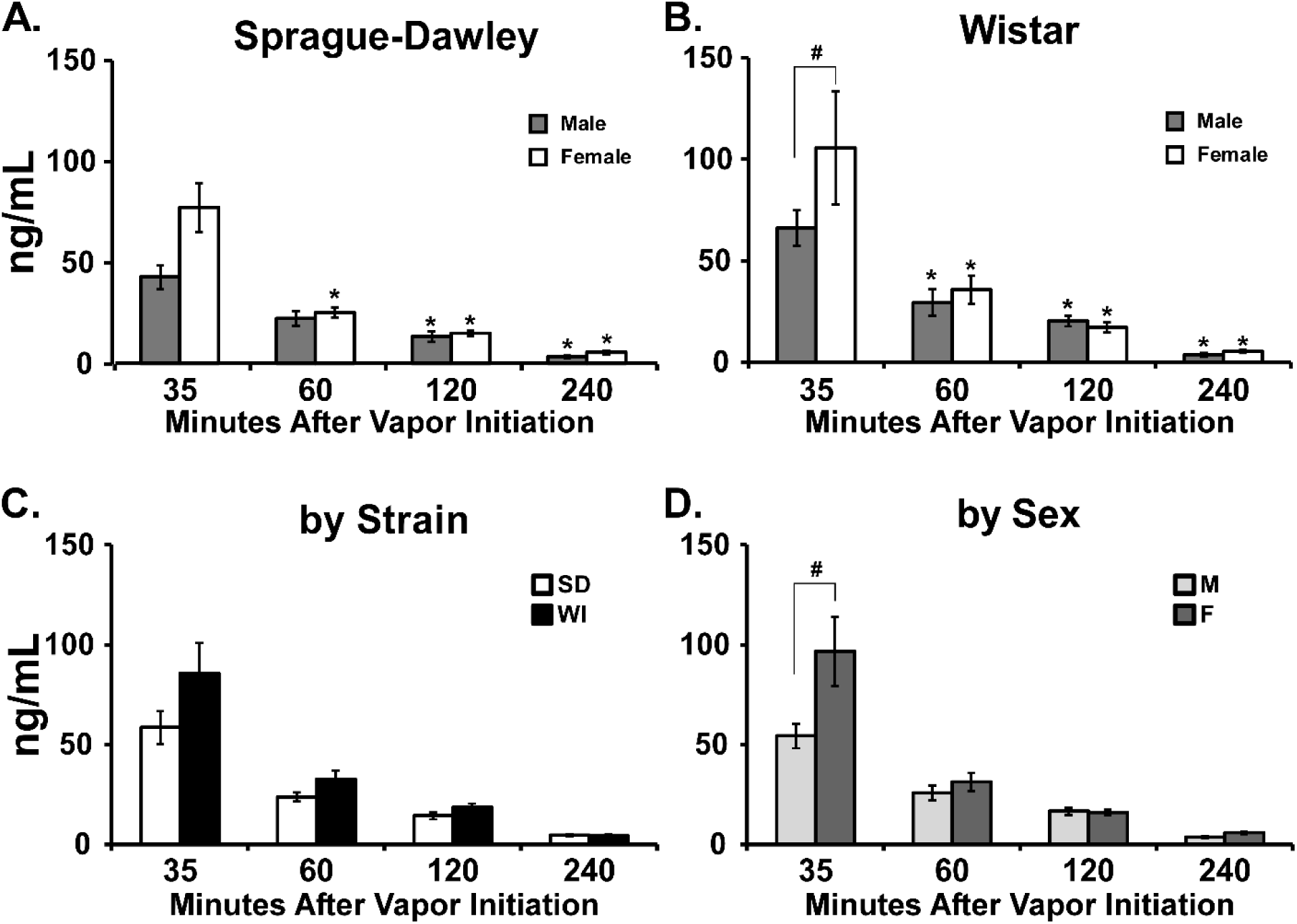
Mean (±SEM) plasma THC concentrations after vapor inhalation of THC (100 mg/mL; 30 minutes) in male and female A) Sprague-Dawley and B) Wistar rats. The data are also presented C) collapsed across sex, by strain (All timepoints differ significantly within SD and WI groups, save 60 and 120 minutes within the WI group do not differ from each other); and D) collapsed across strain, by sex (All timepoints differ significantly from each other within male and female groups). In all panels, a significant difference from the 35 minute timepoint is indicated with * and significant differences between groups with #. The final two timepoints were missing for one WI male and one WI female rat, and the final timepoint was missing for one female SD and one female WI rat in this study.

Insufficient numbers of catheters for the female rats in each group remained patent through the injection study, thus the comparison of i.p. data with the inhalation data focuses only on the male rats (**Figure 8**). The initial mixed-effects analyses confirmed a significant effect of Time after Initiation or Injection on plasma THC levels within the Sprague-Dawley [F (3, 30) = 23.89, P<0.001] and Wistar groups [F (3, 27) = 37.99, P<0.001]. No significant effects of Route of administration or of the interaction of Route with Time were confirmed within either group. A followup analysis was conducted with the males of both strains included in a single group (**Figure 8C**). The mixed-effects analysis again confirmed a significant effect of Time after Initiation or Injection [F (3, 33) = 58.45, P<0.001], but not of Route or the interaction of factors, on plasma THC.

**Figure 8:**
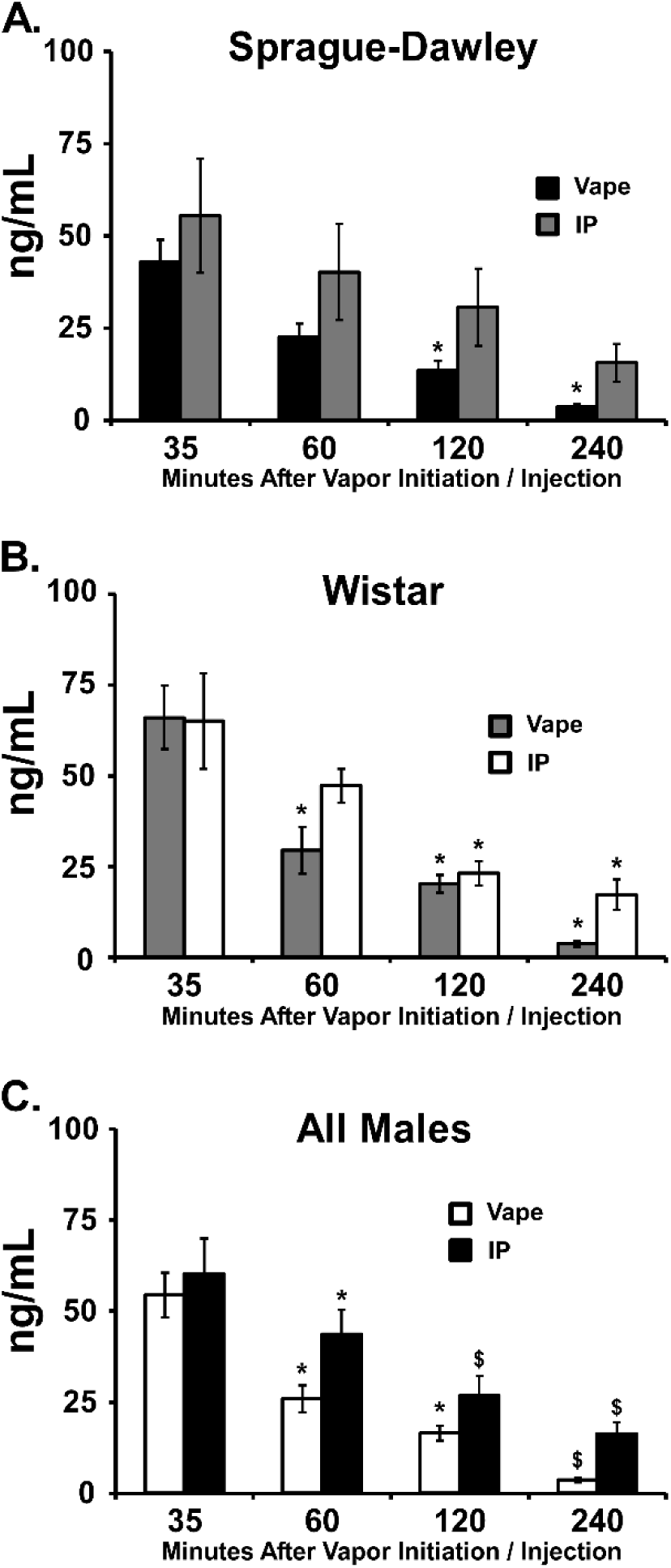
Mean plasma THC concentrations for A) Sprague-Dawley males, B) Wistar males or C) all males after vapor inhalationof THC (100 mg/mL) for 30 minutes or injection with THC (10 mg/kg, i.p.). N=6 per group save that the final timepoint was missing for one WI male in the IP study and the final two timepoints were missing for the same individual in the vapor inhalation study. A significant difference from the 35-minute sample is indicated with * and a difference from both 35-and 60-minute smaples with $.

## 4. Discussion

The present study reports a curious inability of the CB_1_ inverse agonist/antagonists SR141716 or AM-251 to prevent the hypothermia caused by Δ^9^-tetrahydrocannabinol (THC) when delivered by the inhalation route using an e-cigarette vapor system. The antagonists are active at the doses administered, i.e., against the thermoregulatory effects of parenteral injection of THC, and SR141716 attenuated the anti-nociceptive effects of vapor inhalation of THC. Nevertheless, the impact of SR141716 pretreatment on *hypothermia* consequent to THC inhalation is limited to moderately accelerating the return of body temperature to the normal range. We have shown in multiple investigations that the inhalation of vapor produced by dissolving THC in the propylene glycol (PG) vehicle of an e-cigarette device, reduces the body temperature of rats and increases tail-withdrawal latency in a nociception assay (Javadi-Paydar et al., 2019b; Javadi-Paydar et al., 2018; Nguyen et al., 2016b; Nguyen et al., 2019; Nguyen et al., 2018; Taffe et al., 2020). Therefore, this route of administration produces results in rats that are qualitatively similar to effects of THC reported previously in laboratory rodents following parenteral injection (Taffe et al., 2014; Wiley and Burston, 2014; Wiley et al., 2007) and inhalation using different methods (Lichtman et al., 2001; Marshell et al., 2014). Hypothermic effects are observed in male and female Wistar rats and male Sprague-Dawley rats, but are not observed when the rats inhale vapor produced by the PG vehicle alone. Effects on body temperature are also observed in both adolescent and adult rats of both sexes and repeated (2-3 x daily) exposure induces a partial tolerance (Nguyen et al., 2020; Nguyen et al., 2018). Any differences are mostly in the time-course of effects, which is probably attributable to the depot effects of an i.p. injection relative to other routes such as i.v. injection (**Figure 4**) or inhalation. Thus, the present result for the body temperature response identify a case in which effects of THC do not differ qualitatively but the lack of dependence of the initial response on CB_1_ receptor activation appears to be unique to this route of administration.

Positive control experiments within this set of studies confirm that SR141716 and AM-251 (See **Figures 3** and **Supplemental Figure S4**) do produce blockade and reversal of the hypothermia produced by i.p. injection of THC in our model, as has been previously reported for SR141716 (Nguyen et al., 2016b; Taffe et al., 2014), and that produced by i.v. injection as reported herein (**Figure 4**). Numerous prior studies of the hypothermia produced by THC when administered by i.p. injection also show that the body temperature effects can be blocked with pre-treatment of the animals with CB_1_ receptor inverse agonist/antagonists, such as SR141716 or AM-251 (Nava et al., 2000; Nguyen et al., 2016b; Schindler et al., 2017; Taffe et al., 2020). An identical result was found in male rats injected with THC, i.p., in our prior report (Nguyen et al., 2016b). In the present studies, we show further that SR141716 arrests the onset of hypothermia when administered 45 min after i.p. injection, and reverses a substantially developed hypothermia when administered 90 after i.p. injection, of THC. The pharmacokinetic study confirmed that minimal differences in plasma THC are found across these vapor conditions and the same i.p. injection dose at these relevant timepoints (**Figure 8**). Furthermore, the SR141716 dose of 4 mg/kg was sufficient to prevent the hypothermia caused by THC in male and female rats when administered 15 min prior to THC, i.v., and increasing the SR141716 dose to 10 mg/kg in male Wistar rats did not qualitatively alter the failure to arrest the initial hypothermia. Thus it is unlikely that SR141716 is ineffectual to oppose the effects of inhaled THC because of a significant difference in overall, systemic THC dose. Interestingly a prior study of the effects of cannabis smoke inhalation in ICR mice reported no hypothermic effects compared with the placebo smoke condition (Lichtman et al., 2001) which contrasted with a finding in Swiss Webster mice using a THC-infused paper combustion approach (Marshell et al., 2014). As yet unknown strain differences may be critical to the different outcomes in mice, or it could be that subtle methodological differences affect nociception and temperature differently.

This study also confirmed that repeated, twice daily inhalation of THC (100 mg/mL) vapor induces tolerance to hypothermia in adult female rats. Significant tolerance was observed only following the 7^th^ session of a twice daily regimen in adult female rats, consistent with our prior report in which rectal temperature was sampled only after the 1^st^ and 7^th^ sessions (Nguyen et al., 2018) and with our report of a similar timecourse of tolerance development in *adolescent* female rats (Nguyen et al., 2020). We have previously shown that no substantial thermoregulatory tolerance is observed when THC inhalation sessions are spaced by at least 7 days (Javadi-Paydar et al., 2017; Nguyen et al., 2016b) and this was confirmed more directly here since the repeated-exposure animals developed no additional tolerance to intermittent THC challenges (see **Supplemental Figure S3**). Despite this tolerance, presumably mediated at least in part by a down regulation of CB1 mediated signaling, the SR141716 pre-treatment produced a similar result as in the other groups, i.e. no alteration in the initial temperature decrement but an accelerated normalization, post-inhalation. There was a partial attenuation of the initial hypothermia produced by inhaled THC followed by an accelerated return to normative temperature when AM-251 was administered, this can be seen most clearly in **Supplemental Figure 1C**. There was no effect of the CB_2_ receptor antagonist/inverse agonist compound AM-630 on the hypothermic response to inhaled THC, confirming the expected selectivity for the CB_1_ receptor over the CB_2_ receptor. A complete blockade of the effect of i.p. injected THC was, however, produced by SR141716 or AM-251 (**Figure 5**), although interpretation of the effects of AM-251 are somewhat complex because it lowered the body temperature of female rats by itself. Relatedly, a delay in the ability of SR141716 to counter the effects of injected THC, perhaps due to a decrease associated with the SR141716 itself, have occasionally been reported (Fernandez-Ruiz et al., 1997; Taffe et al., 2015).

It is uncertain at present why the two CB_1_ antagonist/inverse agonist compounds did not block the initial hypothermia following inhalation. When administered in these doses, the hypothermia caused by 10 mg/kg THC, i.p., was completely blocked by SR141716 or AM-251 as would be predicted from precedent literature. The nadir of the body temperature observed after inhalation was similar to the nadir reached after i.p. injection, which also suggests a similar effective THC dose in the central nervous system, consistent with the results of the pharmacokinetic study. Importantly, the effects of SR141716 on THC vapor induced anti-nociception were consistent for 90 minutes, starting from immediately post-session. This difference with the thermoregulatory response may suggest differences in THC penetration of hypothalamic regions over spinal-medullary circuitry (Advokat and Burton, 1987; Fields et al., 1983; Fitton and Pertwee, 1982), but it further emphasizes that simple pharmacokinetics do not explain the curious outcome for hypothermia.

Critically, this phenomenon extended in the present study from male Sprague-Dawley and Wistar rats which had not been exposed to repeated-inhalation conditions on sequential days to a generalization and replication of the effect in the female Wistar rats made tolerant by a prior interval of repeated twice-daily THC inhalation. The difference in SR141716 or AM-251 effect across routes of THC administration may be due to the rapidity with which THC enters the brain during inhalation compared with the antagonist brain entry after injection, however this would require further investigation to confirm. The fact that SR141716 blocked any further temperature reduction when administered 45 min after THC, i.p. and almost immediately reversed hypothermia when administered 90 minutes after THC, i.p., argues against an interpretation based on pharmacological competition, however.

There is little doubt that the hypothermia associated with THC inhalation in this model is caused by pharmacological effects of THC. There is never any effect of the PG vehicle inhalation condition on body temperature, ruling out possible hypoxic effects or pharmacological actions of the PG vapor vehicle. The observed *in vivo* effects are related to THC concentration in the vehicle as well as the duration of exposure at a fixed concentration (Javadi-Paydar et al., 2018; Nguyen et al., 2016b; Taffe et al., 2020). It is probable, therefore, that the hypothermia induced by the inhalation of THC is not primarily *initiated* by CB_1_ receptor activity. The CB_1_ receptor does appear to be involved in the *maintenance* of the response in the ∼90 minutes after the nadir is reached and temperature recovery begins to occur. It remains unclear what pharmacological mechanisms might underlie the initiation of hypothermia. Additional studies have shown no ability of the 5-HT1a antagonist WAY 100,635 to block hypothermia induced by THC vapor in male Wistar rats (Javadi-Paydar et al., 2019a) and that the nociceptin/orphanin FQ peptide (NOP) receptor antagonist JTC-801 (1 mg/kg, i.p.) has no effect on hypothermia induced by 20 min inhalation of THC (200 mg/mL) in male Wistar rats (not shown) as predicted by (Rawls et al., 2007). It remains possible that multiple systems are involved in an approximately substitutable manner and that poly-antagonist pharmacology would be required to resolve the mechanism satisfactorily.

## Abbreviations

PG: propylene glycol;
SR141716 (SR): 5-(4-Chlorophenyl)-1-(2,4-dichloro-phenyl)-4-methyl-*N*-(piperidin-1-yl)-1*H*-pyrazole-3-carboxamide;
AM-251: *N*-(Piperidin-1-yl)-5-(4-iodophenyl)-1-(2,4-dichlorophenyl)-4-methyl-1*H*-pyrazole-3-carboxamide;
AM-630: 6-Iodo-2-methyl-1-[2-(4-morpholinyl)ethyl]-1*H*-indol-3-yl](4-methoxyphenyl)methanone;
THC: Δ^9^-tetrahydrocannabinol;

## Author Contributions

MAT and JDN designed the studies, with refinements contributed by KMC, YG, TMK and SAV. JDN, KMC, YG, TMK and SAV performed the research and conducted initial data analysis. MAT conducted statistical analysis of data, created figures and wrote the major drafts of the paper, assisted by JDN. All authors approved of the submitted version of the manuscript.

## Acknowledgements

This work was supported by USPHS grants R01 DA024105 (Taffe, PI), R44 DA041967 (Cole, PI) and R01 DA035482 (Taffe, PI). The National Institutes of Health / NIDA had no direct influence on the design, conduct, analysis or decision to publication of the findings. La Jolla Alcohol Research, Inc. provided funding through a subcontract of R44 DA041967 and donated some necessary equipment and technical support. LJARI did not influence the study designs, the data analysis or the decision to publish findings. Portions of these data were reported in pre-print form in Nguyen et al 2017 (version 1) https://www.biorxiv.org/content/10.1101/172759v1.full (Nguyen et al., 2017).

## Competing Interest

SAV has consulted for LJARI.

## Notes

### Competing Interest Statement

Author SAV has consulted for La Jolla Alcohol Research, Inc., the vendor that supplies our vapor inhalation equipment.

