## Supplemental Materials for "Explication of CB_1_ receptor contributions to the hypothermic effects of Δ9-tetrahydrocannabinol (THC) when delivered by vapor inhalation or parenteral injection in rats"

### S2. Supplemental Methods

S2.1 Supplemental Data/Analysis

*S2.1.1 Supplemental Data/Analysis for Experiment 2 (90 minute pre-teatment with* SR141716 *in male Sprague-Dawley rats) and Experiment 5 (Antagonist Challenge After Repeated THC Vapor Inhalation)*

The main analysis focused on 30 minute averages of the temperature response throughout, but as noted in the Methods, the collection was every 5 minutes. **Figure S1** presents the data that were summarized in Figure 2A,B, and the THC inhalation conditions from the data presented in Figure 5C,D, by 5 minute samples. The data presented in **Figure S2** describe the temperature responses for the PG conditions, and all of the activity data, for the 90 minute pre-treatment study in Experiment 2.

*S2.1.2 Supplemental Data/Analysis for Experiment 5 (Antagonist Challenge After Repeated THC Vapor Inhalation)*

As noted in the main report, these rats received four days of twice-daily inhalation of the PG vehicle then five days of twice daily inhalation of THC (100 mg/mL) during the following week; temperature data are reported for the first session of the day only. Additional THC-inhalation sessions were conducted one and two weeks after the final session of the repeated-THC week and then antagonist studies were initiated 20 days later. The data presented in **Figure S3** show the baseline temperature and the temperature at the 60 minute post-inhalation time point for all of the studies where THC vapor was inhaled with no other treatment or with the vehicle injected, i.p.

S2.2. Supplemental Data Analysis

Data analysis is generally as described in the main Methods with respect to ANOVA / mixed-effects, the factors used and the post-hoc strategy. The 5-minute temperature data are presented for qualitative purposes and are not analyzed with inferential statistics. The activity data for the 90-minute pre-treatment experiment has been grouped by inhalation duration, as the activity for both PG and THC inhalation is presented in this Supplement. For the assessment of additional tolerance in Experiment 5, the critical post-hoc strategy for this study was to use a Dunnett test to compare the temperature response on Day 5 of the chronic week with all other observations post-THC to determine if the intermittent scheduling of the post-chronic assessments induced any additional tolerance.

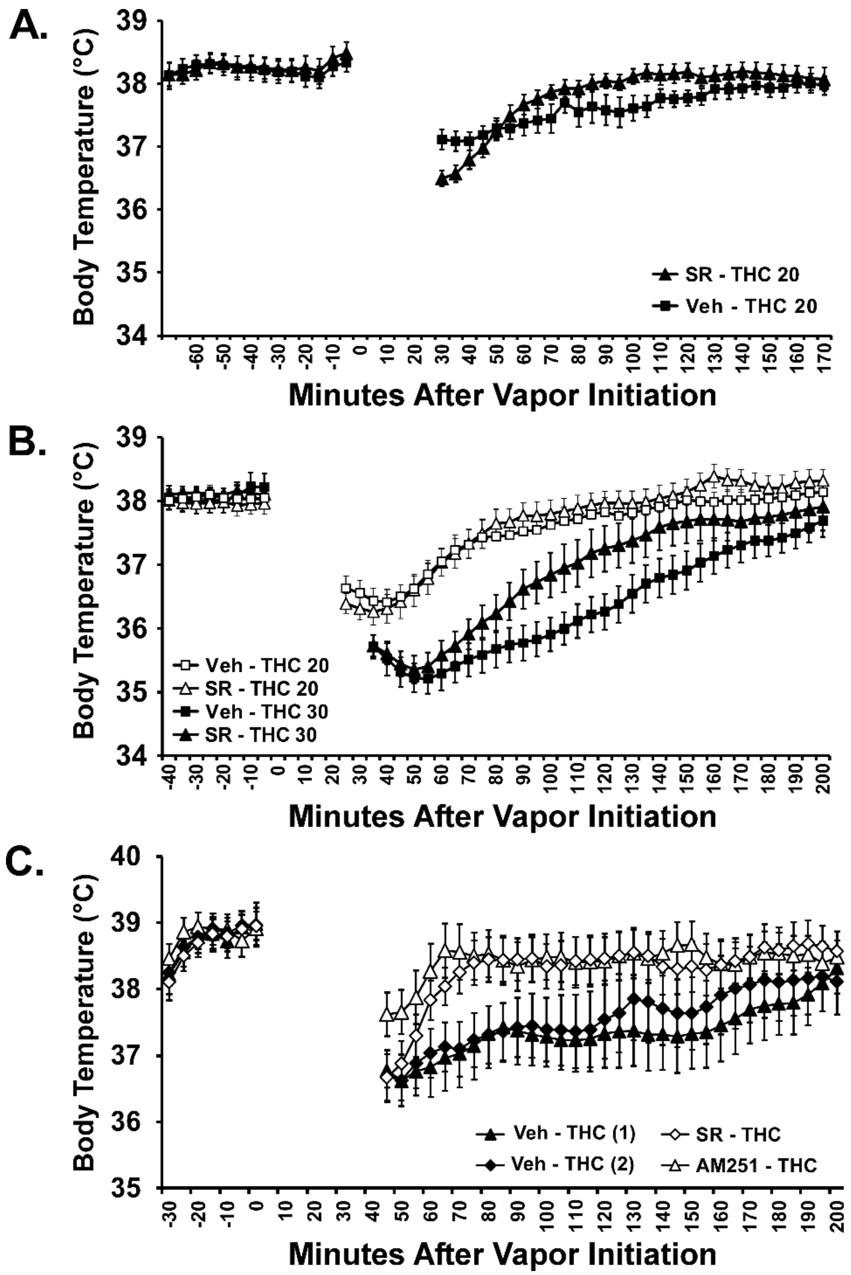


***Figure S1****: Mean (N=8; ±SEM) body temperature of the male Sprague-Dawley rats when SR141716 or the Vehicle was administered 15 minutes (A) or 90 minutes (B) before the inhalation of THC (200 mg/mL) for 20 (THC 20) or 30 (THC 30) minutes. C) Mean (N=8; ±SEM) body temperature of the female Wistar rats when SR, AM-251 or the Vehicle was administered 15 minutes before the inhalation of THC (100 mg/mL) for 30 minutes.*

**S3. Supplemental Results**

*S3.1.Supplemental Results for Experiment 2 (SR141716 in male Sprague-Dawley and Wistar rats) ) and Experiment 5 (Antagonist Challenge After Repeated THC Vapor Inhalation)*

This analysis emphasizes that SR141716 injection prior to THC inhalation for 20 minutes reduced body temperature slightly more than when the vehicle was injected (**Figure S1A, B**). It also emphasizes the similar magnitude of the initial drop in body temperature after 30 minutes of THC inhalation, regardless of the Vehicle/SR141716 pretreatment condition. Furthermore, this analysis makes it clear that AM-251 (4 mg/kg, i.p.) attenuated the initial drop in body temperature (**Figure S1C**); the SR141716 treatment restored temperature to a similar extent as AM-251 in the female group, it just took ~40 minutes longer to do so. The two Veh + THC conditions across those experiments led to identical temperature changes, reinforcing confidence in comparing the effect of the two antagonists without concern about additional tolerance.

As noted in the main report, the inhalation of PG vapor for 20 or 30 minutes had no effect on body temperature (**Figure S2A**) regardless of whether it was preceded by injection of vehicle or SR141716. In the 20 minute THC inhalation condition, there was an apparent increase in activity immediately after vapor inhalation. This was not a timepoint collected in the 30 minute inhalation conditions, thus separate

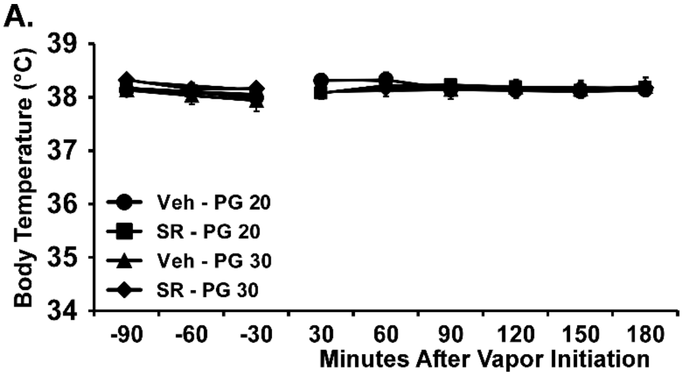


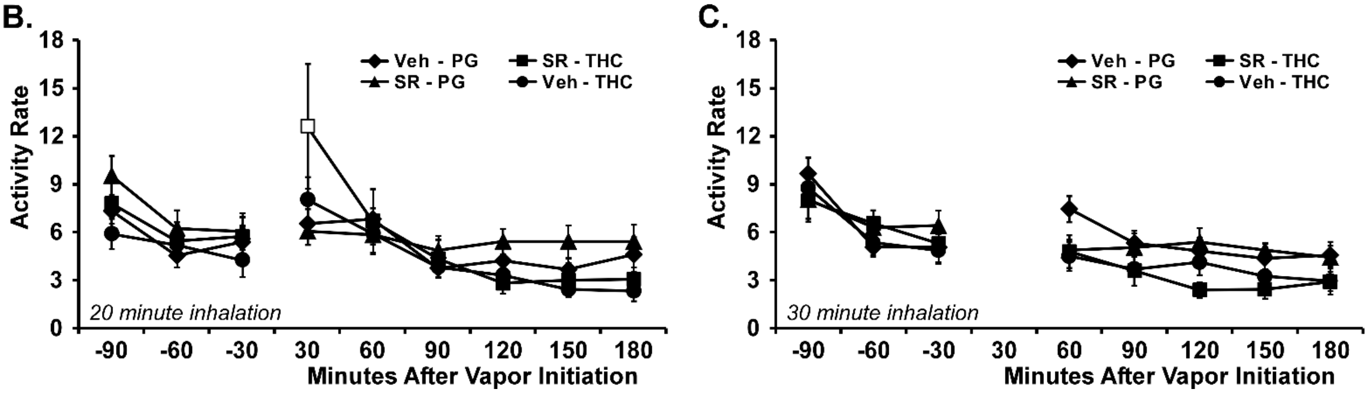


***Figure S2:*** *A) Mean (N=8; ±SEM) temperature of male Sprague-Dawley rats following vapor inhalation of PG for 20 (PG 20) or 30 (PG 30) minutes, initiated 90 minutes after injection with the CB1 antagonist SR141716 (SR; 4 mg/kg, i.p.) or the vehicle (Veh). Mean activity of male Sprague-Dawley (N=8; ±SEM) rats following vapor inhalation of PG or THC for B) 20; or C) 30 minutes. Open symbols reflect a significant difference from the within-treatment baseline and the respective PG inhalation condition.*

analyses focused on all four 20 minute and on all four 30 minute conditions. The three way ANOVA for the 20 minute inhlation conditions confirmed an effect of Time after vapor initiation [F (6, 196) = 8.04;

P<0.0001] and an interaction of Time with Inhalation condition [F (6, 196) = 3.12; P<0.01] on activity rate. There was no significant effect of the pre-treatment condition by itself [F (1, 196) = 3.70; P=0.056], or in interaction with any other factor. The post-hoc test confirmed that activity 30 minutes after the start of inhalation in the SR141716 pretreatment / THC inhalation condition was significantly higher than baseline for the same session and higher than activity at the same time point for the SR/PG condition but not compared with the same timepoint for the Veh/PG or Veh/THC conditions.

The three way ANOVA for the 30 minute inhlation conditions confirmed an effect of Time after vapor initiation [F (5, 168) = 3.83; P<0.005] and of Inhalation condition [F (1, 168) = 21.51; P<0.0001] on activity rate. The post-hoc test did not confirm any significant differences at the 60 minute time point, relative to the within-treatment baseline value, nor any significant differences between treatments at any given time point.

*S3.2. Stability of the THC inhalation-induced hypothermia and injection studies in female Wistar rats*

An analysis of all of the THC-only and THC with Vehicle pre-treatment vapor inhalation studies in this group was conducted to assess the stability of THC-induced hypothermia observed 60 minutes after vapor initiation (i.e., the temperature nadir) across the intermittent challenges conducted subsequent to the repeated THC week. This included follow-up studies of THC 100 mg/mL for 30 minutes conducted one and two weeks after Session 5 and the Vehicle-THC conditions for the pharmacological challenge studies. The ANOVA confirmed a significant effect of Time after vapor initiation [F (1, 7) = 55.18; P=0.0001], of Treatment condition [F (13, 91) = 4.73; P<0.0001] and of the interaction [F (13, 91) = 25.91; P<0.0001]. The Tukey post-hoc test confirmed that significant differences from baseline were observed within each of the THC inhalation sessions. The Dunnett post-hoc confirmed that body temperature 60 minutes after the start of inhalation differed from the 5$^{th}$ THC session in all repeated-PG sessions and the first three repeated-THC sessions but remained unchanged throughout the subsequent studies. In other words, the degree of tolerance induced by the repeated, twice daily exposures was not altered further by up-to-weekly THC exposure.

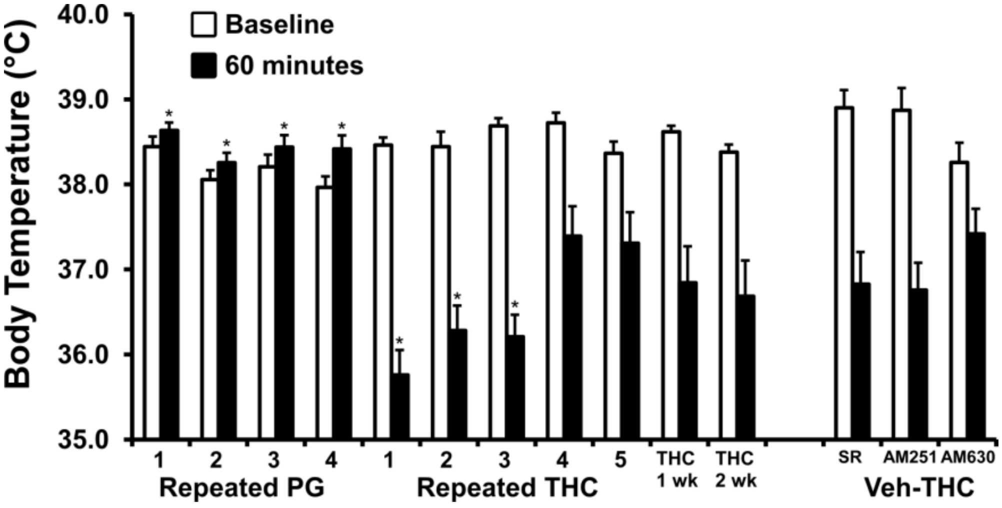


***Figure S3**: Mean (N=8; ±SEM) body temperature of the female Wistar rats before (Base= baseline) and 60 minutes after initiation of vapor inhalation of PG or THC (100 mg/mL) for 30 minutes to compare the hypothermia magnitude across the studies that were conducted in this group. A significant difference from the repeated THC session 5 is indicated with *.*
